# Dissociable effects of prediction and integration during language comprehension: Evidence from a large-scale study using brain potentials

**DOI:** 10.1101/267815

**Authors:** Mante S. Nieuwland, Dale J. Barr, Federica Bartolozzi, Simon Busch-Moreno, Emily Darley, David I. Donaldson, Heather J. Ferguson, Xiao Fu, Evelien Heyselaar, Falk Huettig, E. Matthew Husband, Aine Ito, Nina Kazanina, Vita Kogan, Zdenko Kohút, Eugenia Kulakova, Diane Mézière, Stephen Politzer-Ahles, Guillaume Rousselet, Shirley-Ann Rueschemeyer, Katrien Segaert, Jyrki Tuomainen, Sarah Von Grebmer Zu Wolfsthurn

## Abstract

Composing sentence meaning is easier for predictable words than for unpredictable words. Are predictable words genuinely predicted, or simply more plausible and therefore easier to integrate with sentence context? We addressed this persistent and fundamental question using data from a recent, large-scale (*N* = 334) replication study, by investigating the effects of word predictability and sentence plausibility on the N400, the brain’s electrophysiological index of semantic processing. A spatiotemporally fine-grained mixed effects multiple regression analysis revealed overlapping effects of predictability and plausibility on the N400, albeit with distinct spatiotemporal profiles. Our results challenge the view that the predictability-dependent N400 reflects the effects of *either* prediction *or* integration, and suggest that semantic facilitation of predictable words arises from a cascade of processes that activate and integrate word meaning with context into a sentence-level meaning.

## Introduction

Composing sentence meaning is easier with predictable words than with unpredictable words: for example, ‘bicycle’ is easier to process than ‘elephant’ in “You never forget how to ride a bicycle/an elephant once you’ve learned.” This effect of predictability can be observed in behavioural measures of comprehension such as reading times [1] and on amplitude modulation of the N400 ERP component [2]. The N400 is a negative-going and centro-parietally distributed component of the ERP, which occurs approximately 200-500 ms after word onset and that is strongly associated with lexico-semantic processing [2–3]. Predictable words elicit smaller N400 amplitude than unpredictable words, suggesting facilitation during semantic processing. However, to what extent is such an effect of predictability driven by other relationships between a word and its context, such as the plausibility of the described event? We addressed this issue by re-analysing data from a large-scale (*N*=334) replication study [4]. In a temporally fine-grained analysis, we explored whether predictability, plausibility, and semantic similarity have dissociable effects on word-elicited ERP activity and how these effects unfold over time.

Word predictability is commonly operationalized as ‘cloze probability’, the probability of being used in a non-speeded, offline sentence completion test. The correlation between word predictability and N400 amplitude is well-established [5], with some studies reporting correlations as high as or even higher than *r* = .8 [2, 6–7]. Such results are often considered to demonstrate effects of prediction: people pre-activate a word fully before it appears (as when a specific lexical form can be strongly anticipated in a highly constraining context) or partially (as when some semantic features are activated due to passive spreading of information). Prediction facilitates the semantic activation processes that are initiated when the word appears and are reflected in N400 activity [5].

It is beyond doubt that people can predict the meaning of highly predictable words, and that such predictions ultimately impact the semantic processing of that word. What remains unclear, however, is whether the strong correlations between predictability and N400 amplitude [2,4,6–7] *only* reflect an effect of predictability, or whether they also reflect plausibility of the entire sentence. After all, not only is ‘bicycle’ more predictable than ‘elephant’ given the context “You never forget how to ride a”, it also constitutes a potentially more plausible sentence continuation given one’s long-term knowledge about the world. Plausibility can be established in a norming test in which participants evaluate the plausibility of the described event. Compared to a less plausible word, a more plausible word might be easier to integrate with the context and general world knowledge into a sentence-level interpretation (e.g., [8–9]), regardless of whether word meaning had already been activated before it appeared. If such integration processes are reflected in EEG activity in the N400 time window along with processes of semantic activation [10–11], then the canonical pattern observed for predictability might in part reflect contributions of sentence plausibility.

At the same time, another contribution to the observed patterns might come from a low-level, basic semantic relationship based on simple co-occurrence of words in the context. For example, the word ‘ride’ may occur more often in the context around the word ‘bicycle’ than around ‘elephant’. This low-level semantic relationship called semantic similarity [12] may also contribute to activity in the N400 time window (e.g., [13–15]) and should therefore be acknowledged as a potential factor influencing processing of words in a sentence context (e.g., [15–17]).

Correlations between predictability, plausibility and similarity can make it difficult to establish their effects on processing. Previous studies have tried to avoid this problem by investigating their effects separately. Some studies compare ERPs elicited by equally unpredictable plausible and implausible words [11, 18], whereas others compare ERPs elicited by semantically similar and dissimilar words that are unpredictable and implausible [19–20]. Two recent studies [11,18] report a smaller N400 amplitude for plausible unpredictable words than for anomalous unpredictable words. However, it remains unclear whether these effects purely result from differences in incongruity/plausibility, or whether they (also) reflect differences in semantic similarity. Some studies reported a smaller N400 amplitude for words that are semantically similar to the context, compared to dissimilar words, despite being equally unpredictable and implausible [19–20]. These results suggest that sentence plausibility may have little effect on ERP activity in the N400 time window beyond the combined effects of predictability and semantic similarity. However, Chwilla, Kolk and Vissers [21] demonstrated N400 effects of plausibility for words that were equally unpredictable and equally dissimilar to the context.

These previous studies have only looked at effects of plausibility or similarity on unpredictable, ‘low-cloze’ words, and therefore do not directly address the question of whether or to what extent the well-established, graded relationship between predictability and N400 activity is confounded by other contextual semantic factors. In the current study, we examine the effects of plausibility and similarity across a full range of cloze values. Moreover, we simultaneously model variance associated with predictability, plausibility and semantic similarity as well as lexical variables. By explicitly modelling these sources of variance, we can investigate the effects of one variable while controlling for the others. This approach obviates the need to match conditions on a variable through null hypothesis significance testing, a procedure that is often used but is ineffective and unnecessary [22]. In addition, by modelling activity at individual time samples and EEG channels, we can examine the effects of predictability, plausibility and similarity in terms of both their time course and distribution across the scalp. This contrasts with the ‘traditional’ approach of averaging activity within a longer time window and/or from multiple electrodes (e.g., [2,4,6]. Our use of a spatiotemporally fine-grained analysis is itself not new [15, 23–25], but our study is the first to apply this technique in an attempt to dissociate the effects of predictability, plausibility and semantic similarity.

Our approach can yield new insights into the semantic processes taking place within the first few hundred milliseconds after written word onset. One persistent question in psycholinguistics is whether activity in the traditional N400 time window reflects only the context-dependent activation of semantic information from long-term memory (e.g., [2]), or whether it also reflects integrative processes that compose higher-level sentence meaning from individual word meanings (semantic integration/unification; [8–9]). In other words, it remains unclear whether N400 activity reflects a non-compositional process (activation of meaning) or a compositional process (combining word meanings into a higher-order representation), or perhaps both. Some researchers advocate the latter, ‘multiple-process’ position, and argue that N400 activity does not index a single process but a cascade of semantic activation and integration processes [5, 10, 26–29]. If these processes can be meaningfully studied through their association with predictability and plausibility^1^, respectively, then effects of predictability and plausibility might both be observed in the N400 time window, but effects of predictability would precede and may even be functionally distinct from those of plausibility.

Consistent with this account, a recent study found that effects of prediction appeared earlier than effects of contextual integration [30], and on distinct ERP components (N250 and N400). However, because their participants were instructed to actively predict sentence-final words, the observed ERP patterns may also have reflected task-relevant decision-processes [31], and may not generalize to situations where people do not strategically predict upcoming words (for in-depth discussion, see [32]). The results from studies without such a prediction-task also suggest earlier effects of predictability than of plausibility [11,18], but in these studies the effect of plausibility involved a comparison between plausible and anomalous words, and the contributions of semantic similarity in these studies is unknown.

### The current study

Here, we tested the multi-process hypothesis using data from a large-scale, direct replication attempt of a landmark study on prediction [6]. DeLong et al. capitalized on the phonotactic dependence of the English indefinite articles ‘a/an’ on whether the next word starts with a consonant or vowel. Participants read sentences containing an indefinite article (a/an) followed by a noun. The article-noun pairs were always grammatical but differed in their predictability from sentence context (e.g., “You never forget how to ride <u>a bicycle</u>/<u>an elephant</u> once you’ve learned”). As expected, amplitude of the noun-elicited N400s gradually decreased with increased predictability [2]. Critically, however, DeLong et al. also observed this pattern of results at the preceding articles, which cannot arise from differences in the meaning of ‘a/an’ and therefore does not index integration costs. The article-effect was taken as strong evidence that participants predicted the nouns, including their phonological form (i.e., whether they start with a consonant or vowel), and that the articles that disconfirmed this prediction resulted in processing difficulty (higher N400 amplitude at the article). In a large-scale, direct replication attempt spanning 9 labs [4], our pre-registered analyses did not yield a statistically significant article-effect but successfully replicated the noun-effect.

In the current study, we performed a further (non-pre-registered) analysis to dissociate the effects of prediction and integration. Like previous studies [18–20], we investigated their effects by examining noun-elicited ERP activity as a function of word predictability and plausibility, which we established in offline norms. Improving on previously used methods, we simultaneously modelled variance associated with predictability and plausibility, while also controlling for semantic similarity [12,33], a measure of low-level semantic relatedness between word and context derived from distributional semantics.

If activity in the N400 time window reflects effects of activation and integration [10–11], and if predictability and plausibility reliably correspond to the ease of activation and integration, respectively, then plausibility would have an effect on N400 activity alongside the effect of predictability, although any effects of plausibility would occur later than effects of predictability.

## Methods

Our materials were the 80 sentences in two conditions (expected/unexpected article-noun combination), used by [6]. Participants were native English speaking students from the University of Birmingham, Bristol, Edinburgh, Glasgow, Kent, Oxford, Stirling, York, or volunteers from the participant pool of University College London or Oxford University, who received cash or course credit for taking part in the ERP experiment. Each laboratory aimed to test 40 participants and tested at least 32 participants. For ethical approval and informed consent see [4]. Our data pre-processing was identical to [4], which used a pre-registered procedure (see https://osf.io/eyzaq) that led to the exclusion of 22 participants from the initial group of 356 participants, leaving a sample size of 334 participants. Details about the stimulus materials, participants, EEG recording equipment and settings, experimental procedure and data processing for 22 EEG channels are described [4]. Here, we only describe changes and extensions to the methods from that study. A full list of the materials, including all norming results, is available as Supplementary materials (https://osf.io/47q3s), along with all data, analysis scripts and all supplementary figures. Prior to collecting EEG data, we conducted a predictability (cloze probability) pre-test and a plausibility pre-test.

### Predictability

For the predictability pre-test, we truncated all sentences after the critical indefinite article and asked participants to complete each sentence with the first word or words that came to mind (for details, see [4]). All participants were volunteers from the University of Edinburgh, who did not participate in the ERP experiment or the plausibility tests. We presented two counterbalanced lists of 80 sentences to 30 participants each, such that no participant saw the same sentence context with the expected and the unexpected article. We computed the predictability of each critical noun (the ‘cloze value’) as the percentage of participants who used the word to complete the sentence.

### Plausibility

For the plausibility pre-test, we truncated all sentences after the critical nouns. We presented two counterbalanced lists of 80 sentences to 31 participants each, such that no participant saw the same sentence context with the expected and the unexpected noun. All participants were volunteers from the University of Edinburgh, who did not participate in the predictability pre-tests or the ERP experiment. They were asked to judge “the plausibility of the events described in the sentences”, on a scale from 1 to 7 (from very implausible to very plausible, respectively; other values on the range were shown without a verbal label). We computed a plausibility score for each word as the average of the obtained plausibility ratings over participants. On average, high predictability nouns were rated as plausible (*M* = 5.9, *SD* = 0.48) whereas low predictability nouns were rated as neither plausible nor implausible (*M* = 3.8, *SD* = 1.31).

### Semantic similarity

We measured semantic similarity values with two different techniques. There are emerging developments in the field of distributional semantics and there is ongoing discussion on the suitability of different statistical techniques for extracting and representing the similarity of word meaning text and their relevance for understanding human cognition [33–35]. Traditional ‘counting’ approaches (such as Latent Semantic Analysis, or LSA; [12]) use vectors that count co-occurrences of words in large bodies of text, whereas more recent ‘prediction’ approaches use corpus-trained neural networks to predict words from a set of context words (e.g., word2vec; [35]). Both approaches can successfully model behavioral measures of semantic processing in humans (e.g., semantic priming reaction times), although there are situations where performance of prediction-approaches are superior [33–34].

In a previous version of this manuscript [36], we only used semantic similarity values obtained from word-to-context pairwise LSA [12], based on the General Reading – Up to First Year of College topic space (lsa.colorado.edu), using the maximum factors available. These values correspond to the term-to-term semantic similarity of each target word to its full sentence context. Here, we additionally establish that our key results do not depend on the use of LSA, by repeating our analysis with semantic similarity scores obtained from ‘snaut’ (http://meshugga.ugent.be/snaut), using a word2vec-compatible ‘continuous bag of words’ (CBOW) prediction-model (for details, see [33]). This model was trained on a concatenation of the UKWAC corpus (2 billion words) and a subtitle corpus (385 million words). Values were computed using the English words (300 dimensions, window-size of 6). Where possible, words that did not appear in the corpus were replaced with lemmatized versions^2^.

### Mixed-effects multiple regression

We performed a mixed-effects multiple regression on the single-trial data of [4], obtained with a pre-registered data pre-processing procedure. All segments were resampled to 250 Hz (i.e., one sample every 4 ms). Then, for each sample between −100 to 1000 ms relative to noun onset, and for each channel, we performed a mixed-effects model analysis [37] using the ‘lme4’ package [38] as implemented in R [39]. All continuous predictors (predictability, plausibility, similarity) were z-scored, and we removed random correlations to facilitate model convergence. As expected, our measures of predictability and plausibility were clearly correlated (Spearman’s *r*=0.734, 95% percentile bootstrap confidence interval = [0.726, 0.74]), and more strongly so than predictability and semantic similarity (*r*=0.23 [0.218, 0.241]), and plausibility and semantic similarity (*r*=0.215 [0.203, 0.228]). However, these correlation coefficients are not a principled obstacle to our approach. Variance Inflation Factors (VIF) for all our predictors were below 2.3 (for the additional analyses with interaction terms, VIFs were below 3), which is well below the values deemed problematic due to high multicollinearity [40]. More importantly, multicollinearity is not an issue because the relationship between predictability and plausibility in our items is highly non-linear, which facilitates the disentanglement of their contributions.

In addition, the factor ‘laboratory’ was included as a deviation-coded, categorical fixed effect variable. Although this factor was not of theoretical interest, it was included because our previous pre-registered analysis showed that the overall N400 amplitude differed between laboratories (i.e., a statistically significant main effect of ‘laboratory’; [4]). Because the laboratories did not show significantly different N400 effects of noun-predictability (i.e., no statistically significant interaction between ‘predictability’ and ‘laboratory’), we did not incorporate interaction terms with ‘laboratory’ in the current analyses.

To control for potential N400 effects of lexical frequency and orthographic neighbourhood density [5], we included two additional fixed effects (‘frequency’, using the the logarithmic Zipf scale from the Subtlex-UK database, [41]; and ‘neighbourhood’, using the raw value of Coltheart’s N [42]).

Below, we give the lme4 syntax for the analysis on each time sample and channel (full code for the entire analysis is available on our OSF page). For each analysis, we extracted a coefficient estimate with 95% confidence interval (CI), a *t*-value and *p*-value associated with ‘predictability’, ‘plausibility’ and ‘similarity’. We computed confidence intervals with the ‘Wald’ method, and *p*-values with the normal approximation.

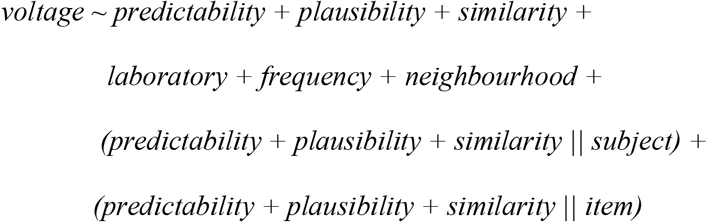

For the purpose of comparison, we also performed the above analysis without the predictors ‘plausibility’ and ‘similarity’, and we also computed ‘traditional’ grand-average ERPs by first averaging trials from either relatively expected and unexpected nouns for each subject and then averaging subject-averages from all subjects per condition. The corresponding Supplementary Figures 4 and 5, respectively, are available on our OSF page.

### Correction for multiple comparisons

We corrected for multiple comparisons using the Benjamini and Hochberg [43] method to control the false discovery rate, the expected proportion of false discoveries amongst the rejected hypotheses. For predictability, plausibility and semantic similarity separately, we applied this correction (implemented in R’s p.adjust) to p-values associated with samples from all electrodes in three time windows of interest (0-200, 200-500, 500-1000 ms). Our main window of interest regarding N400 activity was the 200-500 ms time window, the pre-registered window of analysis in [4], which followed [6]. We applied the correction separately to each window of interest because the false discovery rate procedure, when applied to the entire window, can be too lenient outside the 200-500 ms time window containing large N400 effects [18,24].

### Additional interaction analyses

Our analyses initially did not involve interaction terms, because we did not have a strong a priori theoretical basis to expect the effect size of plausibility or semantic similarity to change with predictability (or vice versa), nor to expect the effect size of plausibility to depend on semantic similarity (or vice versa). However, we subsequently explored potential interactions between our three continuous predictors of interest. No study has yet looked at plausibility effects at medium to high levels of predictability, but effects of either plausibility or similarity have been observed before for unpredictable words [11, 18–21]. In this interaction analysis, we investigate whether the effects of predictability, plausibility and similarity depend on one another, and whether there are effects of plausibility and/or similarity even when accounting for their potential dependence on predictability. We therefore repeated our analysis with the inclusion of all two-way interaction terms between predictability, plausibility and semantic similarity (for both LSA and snaut). To reduce computing time we did not include random slopes for the interaction terms. We applied the same correction for multiple comparisons to the resulting *p*-values as described previously.

## Results

More predictable nouns elicited more positive amplitude (likely indicating a smaller N400 component) than unpredictable nouns within the N400 time window (200-500 ms) across all channels (see Figure 1a, for selected channels; for corresponding scalp distribution plots, see Figure 1d. For waveform plots at all EEG channels, see Supplementary Figure 1 on our OSF page). This effect was statistically significant as early as 200 ms after word onset at multiple channels, and peaked around 330 ms. Following the N400 component, the pattern of activation reversed, such that more predictable nouns elicited a more negative deflection than less predictable items. This post-N400 waveform was statistically significant at frontal and central channels already within the 200-500 ms time window (see also [18]), and appeared stronger and more extended at left- compared to right-hemisphere channels.

**Figure 1.**
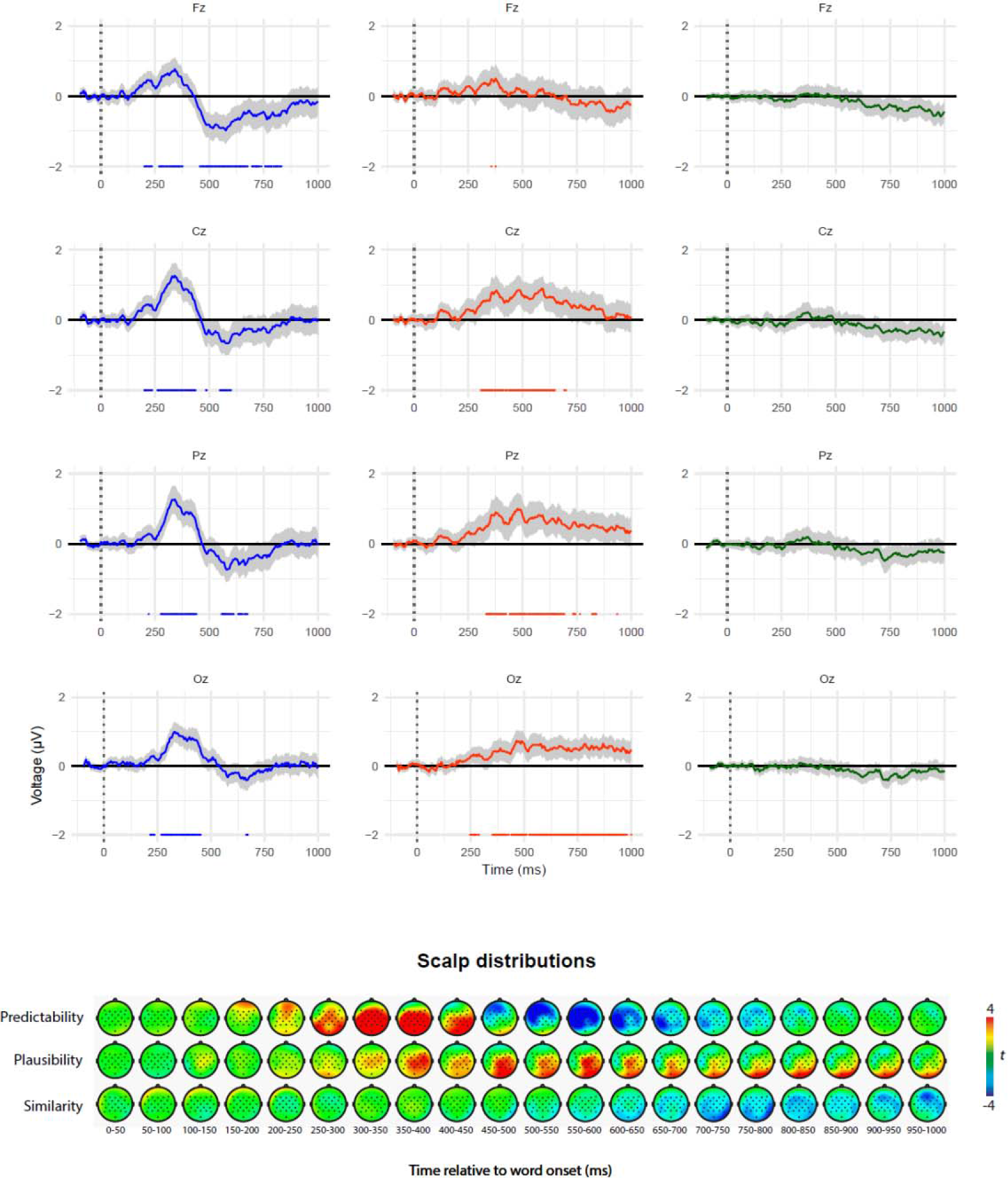
Results from the main analysis. Top graphs: Effect of noun predictability, plausibility, and semantic similarity at 4 midline electrodes. Because we z-transformed these continuous measures, the voltage estimates (coloured lines) and corresponding 95% confidence intervals (grey area) represent the change in voltage, for each time sample and EEG channel, associated with a 1 standard deviation increase. Dots underneath the voltage estimates indicate statistically significant samples after multiple comparisons correction based on the false discovery rate. Lower graphs: Scalp distribution associated with the effects of predictability, plausibility and semantic similarity between the critical noun and the sentence context. The colour scale indexes the average *t*-value within a 50 ms time window relative to word onset.

More plausible nouns elicited more positive amplitude (perhaps indicating a smaller N400 component, but see below) than implausible nouns within the N400 time window (200-500 ms) (see Figure 1b; Supplementary Figure 2). In contrast to the pattern observed for predictability, the effect of plausibility showed a less peaked, more extended time course that continued until about 650 ms after word onset. The effect of plausibility became statistically significant at multiple channels about 350 ms after word onset, thus around the peak effect of predictability, at most channels it peaked or was strongest within the 200-500 ms window, and it was most pronounced over right-hemisphere electrode sites.

We did not observe statistically significant effects of semantic similarity in any of the time windows after multiple comparison correction (Figure 1c; Supplementary Figure 3), but in the 500-1000 ms time window more similar nouns were associated with more negative voltage (see also [36]).

Compared to the effect of predictability in the current analysis, its effect on N400 activity when disregarding plausibility and similarity (Supplementary Figure 5) was overall stronger (as is to be expected when removing a correlated regressor), did not reverse sign at posterior channels like Pz until after 500 ms after word onset, and did not reverse sign at all at right-posterior channels.

In the analysis that included interactions between predictability, plausibility and similarity, none of the interaction terms elicited significant effects after multiple comparison correction (see Figure 2, for selected channels; Supplementary Figures 9a-c), although weak effects were visible. The effect of plausibility became smaller with increasing predictability, and this interaction pattern was strongest at about 500 ms after word onset. The effect of similarity also became smaller with increasing predictability, and this interaction pattern was strongest around the peak of the N400 waveform (~350-375 ms). The effect of similarity became greater with increasing plausibility, and this interaction pattern was strongest right around the peak of the N400 waveform. In this analysis, the main effect of predictability and of similarity remained largely similar to the effect observed in absence of interaction terms, although the effect of similarity now reached statistical significance at two channels around 750 ms after word onset. Importantly, while the main effect of plausibility in this analysis was greatly reduced compared to the effect observed in absence of interaction terms, a clear effect of plausibility was visible and statistically significant just after the N400 peak effect of predictability, in particular at right-hemisphere electrodes.

**Figure 2.**
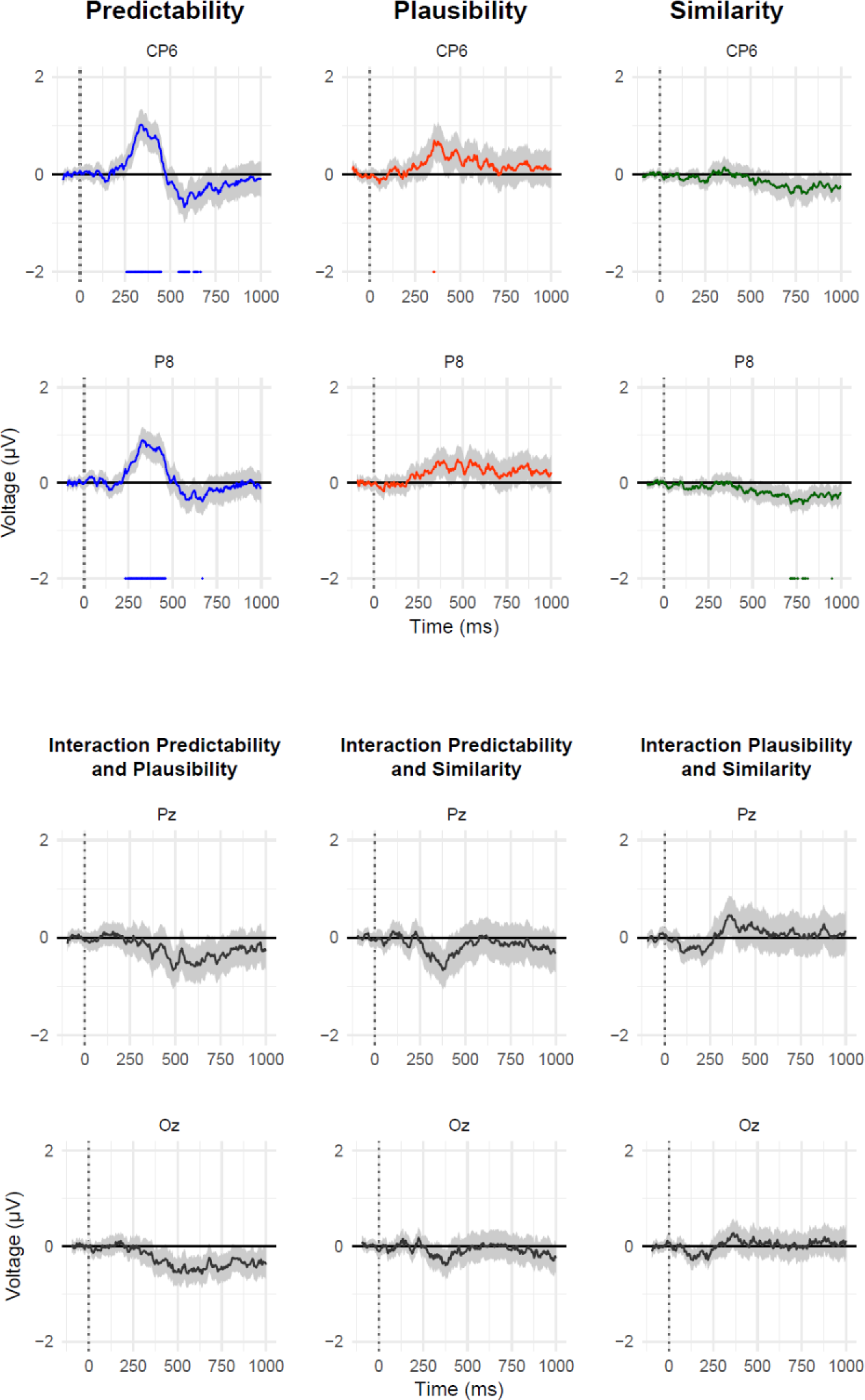
Results from the analysis with interaction terms. Top graphs: Effect of noun predictability, plausibility, and semantic similarity at two right-posterior electrodes. Lower graphs: the voltage estimates (dark grey lines) and corresponding 95% confidence intervals (grey area) corresponding to the 2-way interaction effects between predictability, plausibility and semantic similarity. Negative values indicate that the effect of one variable decreases as the other variable increases.

Finally, our analysis with snaut-based similarity values instead of LSA-values yielded similar results for predictability and plausibility and for their interaction (Supplementary Figures 10-16). Compared to the LSA-analysis, we observed qualitatively different patterns for similarity and associated interaction patterns, but none of these patterns yielded statistically significant effects.

## Discussion

In a reanalysis of data from a large-scale ERP replication study [4], we investigated whether the predictability-dependent N400 [2,6] only reflects the effect of word predictability or also that of sentence plausibility, while controlling for low-level semantic similarity (using different measures of similarity). Unlike previous studies [18–21], we examined the effects of plausibility and similarity across a full range of cloze values, and we simultaneously modelled variance associated with predictability, plausibility and semantic similarity as well as lexical variables. Predictability strongly predicted widespread N400 activity starting as early as 200 ms after word onset, and showed an effect reversal (high predictability eliciting more negative voltage) that started before the end of the N400 window and lasted several hundreds of milliseconds. In contrast, plausibility was associated with a smaller, right-lateralized effect that started only after the effect of predictability reached its peak (around 350 ms after word onset) and that continued until well beyond the classical N400 window. Effects of predictability and plausibility both occurred in the N400 time window, but the former dominated the N400’s rise (i.e., upward flank), while the latter set in at its fall (i.e., downward flank). Semantic similarity did not have a strong effect on N400 activity over and above the effects of predictability and plausibility. Importantly, even when accounting for the possibility that plausibility and similarity have stronger effects for relatively unexpected words, plausibility modulated activity of the N400 after the peak effect of predictability.

If we assume that the association between cloze probability and ERP amplitude reflects an effect of prediction, and that the association between plausibility ratings and ERP amplitude reflects later integration, then our results seem to challenge accounts in which the predictability-dependent N400 reflects the effects of *only* prediction [6] or *only* integration [8]. Instead, they are consistent with a ‘hybrid’, multiple-process view [5,10–11, 27–29], wherein N400 activity reflects a cascade of non-compositional and compositional processes that activate and integrate word meaning within a sentence context. In a recent neurobiological account, Baggio and Hagoort [10] propose that the onset and rising flank of the N400 reflect build-up of current in the temporal cortex when people access word meaning from long-term memory, followed by forward currents to the inferior prefrontal gyrus, where a context representation is generated and maintained. The peak and downward flank of the N400 reflect the moment when re-injection of currents back to the temporal cortex dominates activity as people integrate word meaning with a context representation held active in prefrontal cortex. Our results are broadly compatible with this proposal in terms of the observed time course of prediction and integration effects, although our results are inconclusive with regard to the assumed neural generators. Interestingly, our results do suggest a stronger role for the right-hemisphere in integrative processing, as effects of plausibility were strongest at right-hemisphere electrodes, in accordance with previous literature [5].

Our results obtained in the post-N400 time window (500-1000 ms) inform another current debate, namely on the processing consequences of words that disconfirm a strong prediction. Van Petten and Luka [44] argued for a processing distinction between plausible unexpected words (prediction mismatch) and implausible unexpected words (plausibility violations). The former elicit a left-frontal positive ERP effect, whereas the latter elicit a parietal positive ERP effect [18, 45]. Our study also showed a left-frontal positive ERP effect of prediction mismatch, but the effect was more short-lived (500-750 ms) than is typically reported [44]. We also obtained evidence for a small, late positive ERP effect of semantic similarity, not of plausibility. Therefore, the post-N400 positive ERP effect of prediction mismatch does not reflect effects of prediction and plausibility only. With the caveat that our study did not contain highly implausible semantic anomalies, our results also raise the possibility that positive ERP effects of plausibility violations [44] are at least in part due to low semantic similarity. This further highlights the need to simultaneously model relevant contextual semantic factors to establish the effect of a measure like (im)plausibility.

Our results thus demonstrate a more general point, namely that perhaps any ERP component, and especially those extending over hundreds of milliseconds like the N400 or the post-N400 positivity, is likely to reflect the combined activity of multiple subcomponents that are associated with related yet distinct cognitive processes [29, 46]. Our results could also suggest a reason why N400 effects of contextual congruency sometimes last up to 800 ms after word onset even in written studies [47–48]. Our results suggest that such ‘extended N400 effects’ can occur as the processing consequences of sentence plausibility and may reflect continued efforts to integrate a word with its context (see also [49]). Such effects may be difficult to observe when an opposite ERP effect is simultaneously elicited by another aspect of the stimuli, such as prediction mismatch, but not explicitly modelled in the analysis.

Further research should establish the replicability and generalizability of our results. All our sentences elicited a strong expectation for a given noun [6] and contained indefinite articles that were consistent or inconsistent with that noun (although people may not always use inconsistent articles to revise their prediction, see [4]). In sentences that generate weak or no predictions, integration processes may contribute more strongly to N400. Furthermore, differences between the onset of prediction and integration effects may be exacerbated during spoken language comprehension, as listeners may only need an initial phoneme to disconfirm a prediction [50], well before a lexical meaning is available for contextual integration.

Like previous research [18–20], we investigated semantic activation and integration processes during sentence comprehension by examining the online effects of word predictability and sentence plausibility, which are obtained from separate offline rating tests. However, we acknowledge that the precise relationship between these online processes and offline measures is unknown. It is possible that the observed effects of predictability and plausibility reflect integration processes alone, and that the used predictability and plausibility measures are both noisy estimators of a single underlying function. However, we think this is implausible given the extant literature on the N400 ERP and semantic activation [5]. Similarly, it is possible that both observed effects reflect activation processes alone. However, we think this is implausible given the duration of the observed plausibility effects [5]. In addition, both these accounts are hard to reconcile with the observed changes in effect patterns over time. These issues highlight the need for a detailed and mechanistic account of the transition from activation to integration [27] and, if possible, more direct measures of these assumed processes. Our results suggest that computational models designed to capture context-effects on N400 activity (e.g., [51–52]) and theories of meaning composition more generally, must capture how contributions of context to word-elicited semantic processing can change over time.

In sum, the results of our large-scale study challenge the view that the predictability-dependent N400 reflects the effects of either prediction or integration. Our results suggest that semantic facilitation of predictable words, as reflected in N400 activity, arises from a cascade of processes that activate and integrate word meaning with context into a sentence-level meaning.

## Author Contributions

M. S. Nieuwland developed the study concept. Testing and data collection were performed by E. Heyselaar, E. Darley., S. Von Grebmer Zu Wolfsthurn, F. Bartolozzi, V. Kogan, A. Ito, S. Busch-Moreno, X. Fu., E. Kulakova, S. Politzer-Ahles, and Z. Kohút, under the supervision of M.S. Nieuwland, K. Segaert, N. Kazanina, G. Rousselet, H.J. Ferguson, J. Tuomainen, E. M. Husband, D.I. Donaldson, and S. Rueschemeyer. M. S. Nieuwland performed the data analysis and interpretation in consultation with D. J. Barr, N. Kazanina, G. Rousselet and S. Politzer-Ahles. M. S. Nieuwland drafted the manuscript, and D. J. Barr, F. Huettig, K. Segaert, A. Ito, N. Kazanina, S. Politzer-Ahles, H.J. Ferguson, G. Rousselet, J. Tuomainen, E. M. Husband, D. I. Donaldson, and S. Rueschemeyer provided commentary. All authors approved the final version of the manuscript for submission.

## Footnotes

1 We follow previous work in using these measures as a means to tap into activation and integration processes [e.g., 18–21], with the caveat that their effects may not directly reflect activation and integration processes--i.e., an association between N400 amplitude and cloze probability may not directly reflect active prediction of a stimulus, and an association between N400 and plausibility rating may not directly reflect later integration processes. That is to say, effects of cloze probability may also be driven partly by integration processes and effects of rated plausibility may also be driven partly by activation processes. But if these measures reflect precisely the same processes, then they should not elicit clearly distinguishable effects in neural activity.

2 We thank Pawel Mandera for his generous help with obtaining semantic similarity scores from snaut.

## References

[1] Clifton, C., Jr., Staub, A., & Rayner, K. (2007). Eye movements in reading words and sentences. In R. Van Gompel, M. Fisher, W. Murray, and R. L. Hill (Eds.) Eye movement research: A window on mind and brain. Oxford: Elsevier Ltd. Pp. 341–372.

[2] Kutas, M., & Hillyard, S. A. (1984). Brain potentials during reading reflect word expectancy and semantic association. Nature, 307(5947), 161–163.

[3] Kutas, M., & Hillyard, S. A. (1980). Reading senseless sentences: brain potentials reflect semantic incongruity. Science, 207(4427), 203–205.

[4] Nieuwland, M. S., Politzer-Ahles, S., Heyselaar, E., Segaert, K., Darley, E., Kazanina, N., Von Grebmer Zu Wolfsthurn, S., Bartolozzi, F., Kogan, V., Ito, A., Mézière, D., Barr,D. J., Rousselet, G., Ferguson, H. J., Busch-Moreno, S., Fu, X., Tuomainen, J., Kulakova, E., Husband, E. M., Donaldson, D. I., Kohút, Z., Rueschemeyer, S.-A., & Huettig, F. (2018). Large-scale replication study reveals a limit on probabilistic prediction in language comprehension. eLife, 7: e33468. doi: 10.7554/eLife.33468.

[5] Kutas, M., & Federmeier, K. D. (2011). Thirty years and counting: finding meaning in the N400 component of the event-related brain potential (ERP). Annual Reviews of Psychology, 62, 621–647. doi: 10.1146/annurev.psych.093008.131123

[6] DeLong, K. A., Urbach, T. P., & Kutas, M. (2005). Probabilistic word pre-activation during language comprehension inferred from electrical brain activity. Nature Neuroscience, 8(8), 1117–1121. doi: 10.1038/nn1504

[7] Wlotko, E. W., & Federmeier, K. D. (2012). So that’s what you meant! Event-related potentials reveal multiple aspects of context use during construction of message-level meaning. NeuroImage, 62(1), 356–366.

[8] Hagoort, P., Hald, L., Bastiaansen, M., & Petersson, K. M. (2004). Integration of word meaning and world knowledge in language comprehension. Science, 304(5669), 438–441. doi: 10.1126/science.1095455

[9] Van Berkum, J. J. A, Hagoort, P., & Brown, C. M. (1999). Semantic integration in sentences and discourse: Evidence from the N400. Journal of Cognitive Neuroscience, 11(6), 657–671.

[10] Baggio, G., & Hagoort, P. (2011). The balance between memory and unification in semantics: A dynamic account of the N400. Language and Cognitive Processes, 26(9), 1338–1367. doi: 10.1080/01690965.2010.542671

[11] Lau, E., Namyst, A., Fogel, A., & Delgado, T. (2016). A direct comparison of N400 effects of predictability and incongruity in adjective-noun combination. Collabra: Psychology, 2(1), 13.

[12] Landauer, T. K., & Dumais, S. T. (1997). A solution to Plato’s problem: The latent semantic analysis theory of acquisition, induction, and representation of knowledge. Psychological Review, 104(2), 211–240. doi: Doi 10.1037//0033-295x.104.2.211

[13] Van Petten, C. (2014). Examining the N400 semantic context effect item-by-item: Relationship to corpus-based measures of word co-occurrence. International Journal of Psychophysiology, 94(3), 407–419.

[14] Van Petten, C., Weckerly, J., McIsaac, H. K., & Kutas, M. (1997). Working memory capacity dissociates lexical and sentential context effects. Psychological Science, 8(3), 238–242.

[15] Frank, S. L., & Willems, R. M. (2017). Word predictability and semantic similarity show distinct patterns of brain activity during language comprehension. Language, Cognition and Neuroscience, 1–12.

[16] Nieuwland, M. S., Ditman, T., & Kuperberg, G. R. (2010). On the incrementality of pragmatic processing: An ERP investigation of informativeness and pragmatic abilities. Journal of Memory and Language, 63(3), 324–346.

[17] Otten, M., & Van Berkum, J. J. (2007). What makes a discourse constraining? Comparing the effects of discourse message and scenario fit on the discourse dependent N400 effect. Brain Research, 1153, 166–177. doi: 10.1016/j.brainres.2007.03.058

[18] DeLong, K. A., Quante, L., & Kutas, M. (2014). Predictability, plausibility, and two late ERP positivities during written sentence comprehension. Neuropsychologia, 61, 150–162. doi: 10.1016/j.neuropsychologia.2014.06.016

[19] Federmeier, K. D., & Kutas, M. (1999). A rose by any other name: Long-term memory structure and sentence processing. Journal of Memory and Language, 41(4), 469–495. doi: DOI 10.1006/jmla.1999.2660

[20] Ito, A., Corley, M., Pickering, M. J., Martin, A. E., & Nieuwland, M. S. (2016). Predicting form and meaning: Evidence from brain potentials. Journal of Memory and Language, 86, 157–171. doi: 10.1016/j.jml.2015.10.007

[21] Chwilla, D. J., Kolk, H. H., & Vissers, C. T. (2007). Immediate integration of novel meanings: N400 support for an embodied view of language comprehension. Brain Research, 1183, 109–123.

[22] Sassenhagen, J., & Alday, P. M. (2016). A common misapplication of statistical inference: Nuisance control with null-hypothesis significance tests. Brain and Language, 162, 42–45. doi: 10.1016/j.bandl.2016.08.001

[23] Broderick, M. P., Anderson, A. J., Di Liberto, G. M., Crosse, M. J., & Lalor, E. C. (2018). Electrophysiological correlates of semantic dissimilarity reflect the comprehension of natural, narrative speech. Current Biology, 28(5), 803–809.

[24] Groppe, D. M., Urbach, T. P., & Kutas, M. (2011). Mass univariate analysis of event-related brain potentials/fields I: A critical tutorial review. Psychophysiology, 48(12), 1711–1725. doi: 10.1111/j.1469-8986.2011.01273.x

[25] Hauk, O., Davis, M. H., Ford, M., Pulvermuller, F., & Marslen-Wilson, W. D. (2006). The time course of visual word recognition as revealed by linear regression analysis of ERP data. Neuroimage, 30(4), 1383–1400. doi: 10.1016/j.neuroimage.2005.11.048

[26] Baggio, G. (2012). Selective alignment of brain responses by task demands during semantic processing. Neuropsychologia, 50(5), 655–665.

[27] Baggio, G. (2018). Meaning in the Brain. MIT Press. 2018. ISBN 9780262038126.

[28] Newman, R. L., Forbes, K., & Connolly, J. F. (2012). Event-Related Potentials and Magnetic Fields Associated with Spoken Word Recognition. In: Spivey M, Joanisse M, McRae K, editors. Cambridge Handbook of Psycholinguistics. Cambridge University Press; New York, NY.

[29] Pylkkanen, L., & Marantz, A. (2003). Tracking the time course of word recognition with MEG. Trends in Cognitive Sciences, 7(5), 187–189.

[30] Brothers, T., Swaab, T. Y., & Traxler, M. J. (2015). Effects of prediction and contextual support on lexical processing: Prediction takes precedence. Cognition, 136, 135–149. doi: 10.1016/j.cognition.2014.10.017

[31] Roehm, D., Bornkessel-Schlesewsky, I., Rosler, F., & Schlesewsky, M. (2007). To predict or not to predict: Influences of task and strategy on the processing of semantic relations. Journal of Cognitive Neuroscience, 19(8), 1259–1274.

[32] Nieuwland, M. S. (in press). Do ‘early’ brain responses reveal word form prediction during language comprehension? A critical review. Neuroscience and Biobehavioral Reviews.

[33] Mandera, P., Keuleers, E., & Brysbaert, M. (2017). Explaining human performance in psycholinguistic tasks with models of semantic similarity based on prediction and counting: A review and empirical validation. Journal of Memory and Language, 92, 57–78.

[34] Baroni, M., Dinu, G., & Kruszewski, G. (2014). Don’t count, predict! A systematic comparison of context-counting vs. context-predicting semantic vectors. In Proceedings of the 52nd Annual Meeting of the Association for Computational Linguistics (Volume 1: Long Papers) (Vol. 1, pp. 238–247).

[35] Mikolov, T., Chen, K., Corrado, G., & Dean, J. (2013). Efficient estimation of word representations in vector space. arXiv preprint arXiv:1301.3781.

[36] Nieuwland, M. S., Barr, D. J., Bartolozzi, F., Busch-Moreno, S., Darley, E., Donaldson, D. I., Ferguson, H. J., Fu, X., Heyselaar, E., Huettig, F., Husband, E. M., Ito, A., Kazanina, N., Kogan, V., Kohút, Z., Kulakova, E., Mézière, D., Politzer-Ahles, S., Rousselet, G., Rueschemeyer, S.-A., Segaert, K., Tuomainen, J., & Von Grebmer Zu Wolfsthurn, S. (2018). Dissociable effects of prediction and integration during language comprehension: Evidence from a large-scale study using brain potentials. bioRxiv, 267815.

[37] Baayen, R. H., Davidson, D. J., & Bates, D. M. (2008). Mixed-effects modeling with crossed random effects for subjects and items. Journal of Memory and Language, 59(4), 390–412. doi: 10.1016/j.jml.2007.12.005

[38] Bates, D., Maechler, M., Bolker, B., & Walker, S. (2014). Fitting Linear Mixed-Effects Models Using lme4. Journal of Statistical Software, 67(1), 1–48. doi: 10.18637/jss.v067.i01.

[39] R Core Team (2018). R: A language and environment for statistical computing. R Foundation for Statistical Computing, Vienna, Austria. URL https://www.R-project.org/.

[40] Zuur, A. F., Ieno, E. N., & Elphick, C. S. (2010). A protocol for data exploration to avoid common statistical problems. Methods in Ecology and Evolution, 1(1), 3–14.

[41] Van Heuven, W.J.B., Mandera, P., Keuleers, E., & Brysbaert, M. (2014). Subtlex-UK: A new and improved word frequency database for British English. Quarterly Journal of Experimental Psychology, 67, 1176–1190.

[42] Medler DA, Binder JR. MCWord: An on-line orthographic database of the English language. 2005. http://www.neuro.mcw.edu/mcword/

[43] Benjamini, Y., & Hochberg, Y. (1995). Controlling the false discovery rate: a practical and powerful approach to multiple testing. Journal of the Royal Statistical Society. Series B (Methodological), 289–300.

[44] Van Petten, C., & Luka, B. J. (2012). Prediction during language comprehension: benefits, costs, and ERP components. International Journal of Psychophysiology, 83(2), 176–190. doi: 10.1016/j.ijpsycho.2011.09.015

[45] Quante, L., Bölte, J., & Zwitserlood, P. (2018). Dissociating predictability, plausibility and possibility of sentence continuations in reading: evidence from late-positivity ERPs. PeerJ, 6, e5717.

[46] Dien, J., Michelson, C. A., & Franklin, M. S. (2010). Separating the visual sentence N400 effect from the P400 sequential expectancy effect: cognitive and neuroanatomical implications. Brain Research, 1355, 126–140. doi: 10.1016/j.brainres.2010.07.099

[47] Van Berkum, J. J., Brown, C. M., Zwitserlood, P., Kooijman, V., & Hagoort, P. (2005). Anticipating upcoming words in discourse: evidence from ERPs and reading times. Journal of Experimental Psychology: Learning, Memory and Cognition, 31(3), 443–467. doi: 10.1037/0278-7393.31.3.443

[48] Kulakova, E., & Nieuwland, M. S. (2016). Pragmatic skills predict online counterfactual comprehension: Evidence from the N400. Cognitive, Affective, & Behavioral Neuroscience, 16(5), 814–824.

[49] Romero-Rivas, C., Corey, J. D., Garcia, X., Thierry, G., Martin, C. D., & Costa, A. (2017). World knowledge and novel information integration during L2 speech comprehension. Bilingualism: Language and Cognition, 20(3), 576–587.

[50] Van Petten, C., Coulson, S., Rubin, S., Plante, E., & Parks, M. (1999). Time course of word identification and semantic integration in spoken language. Journal of Experimental Psychology: Learning Memory and Cognition, 25(2), 394–417. doi: Doi10.1037//0278-7393.25.2.394

[51] Fitz, H., & Chang, F. (2018). Sentence-level ERP effects as error propagation: A neurocomputational model. PsyArxiv.

[52] Rabovsky, M., Hansen, S. S., & McClelland, J. L. (2018). Modelling the N400 brain potential as change in a probabilistic representation of meaning. Nature Human Behaviour, 2(9), 693.

